# Extending the CWM approach to intraspecific trait variation: how to deal with overly optimistic standard tests?

**DOI:** 10.1101/2021.09.09.459685

**Authors:** David Zelený, Kenny Helsen, Yi-Nuo Lee

## Abstract

Community weighted means (CWMs) are widely used to study the relationship between community-level functional traits and environment. For certain null hypotheses, CWM-environment relationships assessed by linear regression or ANOVA and tested by standard parametric tests are prone to inflated Type I error rates. Previous research has found that this problem can be solved by permutation tests (i.e. the max test). A recent extension of the CWM approach allows the inclusion of intraspecific trait variation (ITV) by the separate calculation of fixed, site-specific, and intraspecific CWMs. The question is whether the same Type I error rate inflation exists for the relationship between environment and site-specific or intraspecific CWM. Using simulated and real-world community datasets, we show that site-specific CWM-environment relationships have also inflated Type I error rate, and this rate is negatively related to the relative ITV magnitude. In contrast, for intraspecific CWM-environment relationships, standard parametric tests have the correct Type I error rate, although somewhat reduced statistical power. We introduce an ITV-extended version of the max test, which can solve the inflation problem for site-specific CWM-environment relationships and, without considering ITV, becomes equivalent to the "original" max test used for the CWM approach. We show that this new ITV-extended max test works well across the full possible magnitude of ITV on both simulated and real-world data. Most real datasets probably do not have intraspecific trait variation large enough to alleviate the problem of inflated Type I error rate, and published studies possibly report overly optimistic significance results.

## Introduction

According to community assembly theory, which species will occur in a local community partly depends on the result of environmental filtering by the prevailing local abiotic conditions (Keddy 1992, Zobel et al. 1998). More recently, this environmental filtering is believed to act directly upon species’ functional response traits (Lavorel & Garnier 2002). These traits consist of measurable properties of an individual organism that directly influence its fitness under the prevailing environmental conditions (Violle et al. 2007). The realisation of this link between functional traits and the environment has opened up avenues to uncover the mechanisms behind community assembly, and to predict community responses to environmental change. This has resulted in an ever-increasing number of studies exploring functional trait-environment relationships (e.g. Miller et al. 2019).

At the community level, trait-environment relationships are regularly assessed by calculating community weighted mean trait values (CWMs) (Garnier et al. 2004, Diaz et al. 2007). The resulting CWMs are then usually directly related to different environmental variables using correlation, regression, ANOVA or other general(ized) linear mixed model techniques. We call this the *CWM approach* in this study. A number of alternative methods are also available for assessing trait-environment relationships, including the fourth corner (Legendre et al. 1997, Dray & Legendre 2008, Peres-Neto et al. 2017), species’ niche centroids (SNC; Peres-Neto et al. 2017, ter Braak et al. 2018a), multilevel models (Brown et al. 2014, Jamil et al. 2013, Warton et al. 2015, Miller et al. 2019), bootstrapping approach (Maitner et al. 2023) or double constrained correspondence analysis (ter Braak et al. 2018b).

Traditionally, the CWM approach used fixed species-level trait values (a given species has the same trait value in all occupied sites) and thus ignored intraspecific trait variation (ITV), i.e. variation in trait values among individuals of the same species. This was justified by the assumption that, in most datasets, the amount of ITV is negligible compared to the amount of interspecific trait variation, i.e. variation among species (McGill et al. 2006). However, this assumption has recently been challenged by several studies that found that both within- and among-community ITV is often substantial, at least for plants (Albert et al. 2010, Messier et al. 2010, Siefert et al. 2015, Westerband et al. 2021), while less well understood for other taxa (e.g. Gaudard et al. 2019 for ants, Behm and Kiers 2014 for arbuscular mycorrhizal fungi, or Dawson and Jönsson 2020 for basidiomycetes). ITV has two main sources, namely genetic variation (there can be several genotypes present in the population) and phenotypic plasticity resulting from the response of individuals (or their parents) to environmental heterogeneity (Geber & Griffen 2003). Consequently, researchers are now actively advocating the inclusion of ITV in most community ecological trait research, including in trait-environment relationship tests (Albert et al. 2011).

Specifically for CWM trait-environment relationships, studies have clearly illustrated that results can be biased when using fixed species-level trait values instead of incorporating ITV (Albert et al. 2012, Borgy et al. 2017). This has resulted in an increasing number of studies where authors calculate CWMs using site- or habitat-specific trait values, measured separately for each species in each site or habitat, respectively. Lepš et al. (2011) introduced an extension of the CWM approach that allows the partitioning of the relative contribution of ITV and interspecific trait variation to community-level trait-environment relationships. This approach is based on the realisation that CWMs calculated from fixed species-level trait values ("fixed" CWM, excluding ITV) can vary among communities only if their species composition differs (species turnover). On the contrary, differences in CWMs calculated from site-specific trait values ("site-specific" CWM) can be caused by both species turnover and ITV. The difference between the site-specific CWMs and fixed CWMs then only encompasses the effect of ITV ("intraspecific variability effect" CWM, or "intraspecific" CWM in short). These fixed, site-specific and intraspecific CWMs are subsequently related to environmental variables using either regression or ANOVA, and their explained variation is partitioned between species turnover and ITV. This method has proven popular, and to date, we have identified over 60 published case studies using it. We are not aware of any other published method which would, at the community level, aim to disentangle and quantify the effect of interspecific and intraspecific trait variation.

Several studies recently found that the standard parametric tests in the CWM approach (relating CWM trait values to species occurring in the community to environmental variables of its habitat) using fixed trait values are prone to Type I error inflation. This results in the situation that even CWMs calculated from randomly generated species-level trait values often show significant correlations to environmental variables (Peres-Neto et al. 2017, ter Braak et al. 2018a, Zelený 2018; but see Lepš & de Bello 2023 for an alternative view). Peres-Neto et al. (2017) have shown that the correlation of CWMs and environmental variables is in fact numerically tightly related to the fourth corner method (introduced by Legendre et al. 1997), and that the same solution used for controlling the Type I error rate in the fourth corner (Dray et al. 2008, ter Braak et al. 2012), can be used to control the inflated Type I error rate in the CWM approach. This solution is based on a combination of two permutation tests, one permuting the sample attribute (i.e. the environmental variable) and the other permuting the species attribute (i.e. the trait), into the "max test", by taking the higher (more conservative) P-value of the two permutations (Cormont et al. 2011, ter Braak et al. 2012).

However, it is unclear if this Type I error inflation persists when introducing ITV in the CWM approach following the method of Lepš et al. (2011). To date, none of the studies using this method have tested or tried to correct this potential Type I error inflation. Note, however, that Candeias and Fraterrigo (2020) and Sandel and Low (2019) partly acknowledged and tried to address related Type-I error inflation issues in their studies. Part of the reason why the Type I error problem for fixed CWMs arises is that the calculated CWMs of different sites in a dataset are often not independent since some pairs of sites usually share at least some species, and the trait values of these shared species are identical. An extreme case would be the situation when two sites share all species, and these species have the same relative abundances; CWMs calculated for each of these sites must then be identical. This lack of independence between fixed CWMs reduces the effective degrees of freedom in the analysis of their relationship with environmental variables and causes the Type I error inflation to be negatively dependent on the beta diversity of the species composition dataset (Fig. 3a in Zelený 2018). We expect that this problem will be relaxed for site-specific CWMs because the inclusion of ITV allows species to have different trait values in different sites, thus reducing the dependence among sites. However, although ITV can be substantial, at least for plants, it is often smaller than interspecific trait variation (cf. Messier et al. 2010, Siefert et al. 2015, Westerband et al. 2021). We consequently expect that using site-specific trait values will not completely remove the dependency issue but that the severity of inflation will depend on the magnitude of ITV. Moreover, the currently available max test solution cannot be applied because the vector-based trait permutation for fixed CWMs cannot readily be extended to the site-by-species matrix for site-specific CWMs. For intraspecific CWMs, we do not have enough clues to forecast whether they are or are not affected by inflated Type I error rate.

In this study, we explore 1) whether the CWM approach suffers from inflated Type I error rates when including ITV, by calculating site-specific and intraspecific CWMs, 2) whether this potential inflation depends on the magnitude of ITV, and 3) whether our newly proposed modification of the max test can overcome this potential inflation problem. To explore these questions, we quantified Type-I error rates for simulated community data with varying levels of intra- and interspecific trait variation, and a real-world dataset consisting of four functional leaf traits measured along a wind gradient for cloud forest vegetation in northern Taiwan.

## Materials and Methods

### Community weighted mean approach and extension for intraspecific trait variation

When individual trait-environment relationships are analysed at the community level, three objects are usually involved: a species composition matrix, an environmental variable (vector) and a species trait (vector). Species composition is represented by an *n*-by-*S* matrix **L** = [*l_ij_*], where *n* is the number of sites (rows), *S* is the number of species (columns), and *l_ij_* is the contribution of species *j* to site *i* (where contribution can be expressed as abundance, biomass, cover or another quantitative measure, or as presence-absence). The environmental variable is represented by a *n*-elements-long vector **e** = [*e_i_*], where *e_i_* is the value of the environmental variable for site *i*. The trait is represented by an *S*-elements-long vector **t** = [*t_j_*], where *t_j_* is the trait value of species *j*. Naming conventions follow Peres-Neto et al. (2017), with a few exceptions (explicitly mentioned further in the text) and several extensions.

The CWM approach first translates the species-level vector **t** to a site-level vector **c** = [*c_j_*], by calculating the average trait value for a site across all present species, weighted by each species contribution, as expressed in the matrix **L**. The community weighted mean for site *i* is calculated as 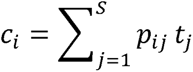, where *p_ij_* is the relative contribution of species *j* in site *i*, and *t*_j_ is the fixed trait value of species j (Garnier et al. 2004, Díaz et al. 2007). Relative contribution *p*_ij_ is calculated by dividing *l*_ij_ by the sum of species contributions in site *i* for which trait values are available, i.e. as 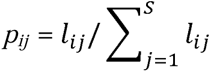. Species with missing trait values should not be included in the calculation of *p_ij_*, so that the sum of relative species contributions in site *i* is always equal to one (Zelený 2018). Next, vector **c** is directly related to the environmental vector **e** by correlation, regression, ANOVA or another method, and the significance of this relationship is often tested.

Extension of the CWM approach to allow the inclusion of ITV is done by distinguishing site-specific and fixed trait values (Lepš et al. 2011). Site-specific trait values for species *j* in site *i* thus become an *n*-by-*S* matrix **T** = [*t_ij_*], where *t_ij_* represents the mean trait value calculated from individuals of species *j* collected within site *i* (the value is missing if the species does not occur at the site or none of its individuals has been measured). The fixed trait values are denoted as a *n*-elements-long vector **t̄** = [*t̄*_j_], where *t̄*_j_, is calculated as the mean of all site-specific trait values (*t_ij_*) of species *j* across all *n* sites in the dataset where that species occurs.

Using the site-specific (**T**) and fixed (**t̄)** trait values, Lepš et al. (2011) calculated site-specific CWMs, which include ITV as 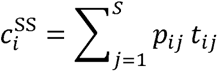, resulting in a *n*-elements-long vector 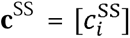, while fixed CWMs, which do not consider ITV, were calculated as 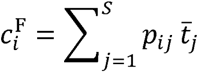, resulting in a *n*-elements-long vector 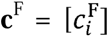, that is essentially *c*_i_ as calculated in the absence of ITV measurements (if we assume that *t̄*_j_ = *t̄*_j_). Finally, the intraspecific variability effect (called intraspecific CWM here) is defined as the difference between the site-specific CWM and the fixed CWM and calculated as 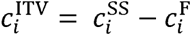, stored in a *n*-elements-long vector 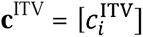. Using the above formulas, the calculation of **c**^ITV^ can be rewritten to 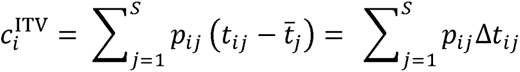, where Δ*t_ij_* = *t_ij_* − *t̄_j_* are site-specific trait values centred by species, represented by a *n*-by-*S* intraspecific trait matrix Δ**T =** [Δ _ij_]. Thus, unlike c^SS^, which quantifies the absolute CWM trait values observed at different sites, c^ITV^ only quantifies the contribution of ITV to site-specific CWMs.

Lepš et al. (2011) pointed out that changes in site-specific CWMs (**c**^SS^) are caused either by species composition turnover (quantified by **c**^F^), changes in species-level trait values, i.e. ITV (quantified by **c**^ITV^), or by both. They proposed a method to partition the effect of these two sources, in which **c**^SS^, **c**^F^ and **c**^ITV^ are separately related to the vector **e** using a general linear model approach. The sum of squares (SS) are then extracted from each model, where SS_specific_ represents the total among-site trait variation explained by the environmental variable (**c**^SS^ ∼ **e**), while SS_fixed_ and SS_intra_ represent the contribution of species turnover (**c**^F^ ∼ **e**) and ITV (**c**^ITV^ ∼ **e**), respectively. If the effects of species turnover and ITV vary independently, SS_specific_ = SS_fixed_ + SS_intra_. Usually, however, the effects of species turnover and ITV covary, either positively (i.e. when **c**^F^ and **c**^ITV^ both have either a positive or a negative response to the environmental variable) or negatively (i.e. when **c**^F^ and **c**^ITV^ respond oppositely to the environmental variable). Lepš et al. (2011) therefore suggested adding a covariation component, calculated as SS_cov_ = SS_specific_ – SS_fixed_ – SS_intra_. This approach has been introduced by Lepš et al. (2011) in their supplementary material, and implemented in the R package *cati* (Taudiere & Violle 2015).

### Type I error inflation for trait-environment relationships in the CWM approach

Previous studies have shown that using the CWM approach without considering ITV (thus using the fixed CWM vector c^F^) to assess the link between environment and traits often results in Type I error inflation. This is explained by the fact that a true link between traits and environment (**t**↔**e**) can only occur if the link between environment and species composition, and between traits and species composition are present (**e**↔**L** and **t**↔**L**, respectively). Type I error inflation occurs when the environment, but not traits, is related to the species composition (i.e. **e**↔**L** and **t**↮**L**, Peres-Neto et al. 2017). The solution to this inflation problem was adopted for the CWM approach by Peres-Neto et al. (2017) from an analogous solution applied in the fourth-corner approach (Legendre et al. 1997; Dray & Legendre 2008). It consists of calculating two permutation tests, one permuting the rows in **L** to test the **e**↔**L** link and one permuting the columns in **L** to test the **t**↔**L** link. Both tests are then combined together into the "max test" by only taking the largest P-value (least significant result) as the test of the **t**↔**e** link (Cormont et al. 2011, ter Braak et al. 2012). An equivalent result is achieved by replacing the row-based permutation of the **L** matrix by permuting vector **e** and relating it to vector **c**^F^ (calculated from not-permuted trait values **t**), and replacing the column-based permutation test of **L** by permuted trait values **t**, and relating the newly resulting vector ^P^c^F^ (where ^P^ stands for permuted) to the not-permuted vector **e** (ESM Fig. S1; Zelený 2018). For convenience, we still refer to these permutation schemes as row- and column-based permutations, respectively.

### Simulated community data with ITV

To assess whether the CWM approach extended for ITV has a correct Type I error rate, we simulated community data that included intra- and interspecific trait variation that was directly structured by a hypothetical environmental gradient. More specifically, we evaluated the potential Type I error inflation for the linear regressions between vectors **c**^F^, **c**^SS^ or **c**^ITV^, on the one hand, and vector **e**, on the other hand. The description of how we generated simulated community data by an R library `simcom` is detailed in ESM Methods S1.

We set the number of species to 50, the number of sites to 25, and each simulation was performed 50 times with the same number of species and sites, resulting in 50 independent sets of **e** and **L**. For each simulation, we generated a matrix of simulated site-specific trait values **T**, in which both interspecific and intraspecific trait variation was completely, positively linked to vector **e**. The *n*-by-*S* matrix **T** was first generated by replacing each non-zero value of *l*_ij_ in site *i* of matrix **L** with the value *e*_i_ (zero values of *l*_ij_ became missing values in **T**). To allow modifying the magnitude of ITV in the simulated matrix **T**, we first calculated the vector of fixed trait values as the means of individual columns of matrix **T**, and calculated the matrix **ΔT** = [Δ*t_ij_*] = [*t_ij_* − *t̄_j_*]. We then introduced coefficient *m* to control the magnitude of simulated ITV, and calculated the matrix of site-specific trait values as 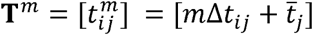. For each simulated vector **e** and matrix **L** we generated a set of site-specific trait matrices **T***^m^* for *m* ranging from 0 to 5 with 0.5 intervals. If *m* = 0 (no ITV) all values of _ij_ in column *j* are identical, and equal to _j_ ; this matrix was reduced into the vector and used as fixed trait values. Increasing *m* increases the magnitude of ITV (for *m* = 1, the values in _ij_ are identical to those calculated from **L** and **e**, as described earlier). Additionally, we also constructed a simulated **T**^m^ matrix in which the trait values were random values drawn from a standard normal distribution, N (0, 1); no random noise was added to *t*_ij_ value in this scenario. The latter scenario (with notation *m* = ∞ used in the following text) represents a situation in which the site-specific trait value of a given species is a random sample from the full pool of potential trait values across all species, unconstrained by any species-specific ITV range. Thus, in total, we have one vector of fixed trait values and 11 site-specific trait matrices **T**^m^ for each simulation. Finally, we rescaled all values in **T** into the range between 0 and 1, and added a small value (generated as a random number from a uniform distribution between -0.1 and 0.1) as a random noise to each *t*_ij_.

### Dependence of the Type I error rate on the magnitude of ITV

To assess the Type I error rates of the linear regressions of **c**^F^, **c**^SS^ or **c**^ITV^ to vector **e,** tested by parametric *F*-tests using our simulated community data, we first cancelled the link between the traits and the species composition by permuting the trait matrices for each separate simulation (ESM Fig. S2). For the fixed trait values, we permuted values within the vector **t̄** to get ^P^**t̄** = [^P^*t̄_j_*] (ESM Fig. S2). For the intraspecific trait values, we calculated the intraspecific trait matrix Δ**T^m^**for each **T^m^** and permuted the values within each column (species) of Δ**T**^m^ to get ^P^Δ**T**^m^ = [^P^Δ*t_ij_*]. Note that these column permutations are only performed across cells where the species is present. Finally, vector ^P^**t̄** and matrix ^P^Δ**T**^m^ were combined into a matrix of permuted site-specific trait values ^P^**T**^m^ = [^P^*t_ij_*] = ^P^Δ*t_ij_* + ^P^*t̄_j_*. Subsequently, ^P^**t̄**, and all ^P^**T**^m^ and ^P^Δ**T**^m^ matrices were combined with the **L** matrix to calculate one ^P^**c**^SS^, 11 ^P^**c**^F^ and 11 ^P^**c**^ITV^ vectors, respectively. Each of these 23 CWM vectors was then regressed against vector **e,** and significance levels were assessed by parametric *F*-tests. We repeated all trait matrix permutations 1000 times, and for each of the 23 trait-environment regressions, we counted the number of significant correlations (p < 0.05) (N_obs_). Since the null hypothesis that the trait is not correlated to the environmental variable is true, because we broke the link between the trait and the species composition by permuting trait values, the expected number of significant correlations (i.e. the Type I error rate) is α (0.05) × 1000 = 50 (N_exp_). The Type I error rate inflation was then quantified using the inflation index I(α) = N_obs_/N_exp_ (Lennon 2000); an index value of 1 indicates no inflation.

Finally, the mean ± SD inflation index across the 50 simulations was plotted against the magnitude of ITV in the simulated data (cf. parameter *m*), separately for ^P^c^SS^, and ^P^c^ITV^. To allow comparison with real datasets, we transformed parameter *m* to "relative ITV index", calculated as the mean variance of ITV (i.e. variance of individual species’ site-specific trait values across all sites, averaged across species) divided by the variance of interspecific trait variation across all species included in the analysis (calculated from the fixed trait values). This ratio is equivalent to the *T_PC/PR_* T-statistic introduced by Violle et al. (2012).

### Introducing a max solution for the ITV-extended CWM approach

The previously described row- and column-based permutation tests used for controlling Type I error inflation in the CWM approach (Peres-Neto et al. 2017, ter Braak et al. 2018a) cannot be directly used for the ITV-extended CWM approach (Lepš et al. 2011). While the row-based permutation test can still be performed by permuting vector **e** (Fig. 1a), it is not clear how the column-based permutation test should permute trait values in the trait matrix **T**, as opposed to vector used in the absence of ITV. Both the species composition matrix (**L**) and the site-specific trait matrix (**T**) usually contain some sites *i* where species *j* is absent (*p_ij_* = 0), and thus the site-specific trait value is not available (*t_ij_* = NA). Permuting columns in **T** (analogously to permuting elements in **t**) would mismatch values in these two matrices, causing some species with non-zero abundances in **L** to be newly paired with missing site-specific trait values in **T** and vice versa. To avoid this problem, we propose a new max test version containing a column-based permutation test that combines separate interspecific trait permutation (on vector ) and intraspecific trait permutation (on matrix Δ**T**). For vector **t̄**, the fixed trait values are directly permuted to obtain ^P^**t̄**. The permutation of values in Δ**T**, however, is done separately for each column (ignoring cells with missing values) to get ^P^Δ**T**. The permuted mean trait values in vector ^P^**t̄** and permuted site-specific trait matrix ^P^Δ**T** are then combined together into a new matrix of permuted site-specific trait values ^P^**T**. This new matrix is then used to calculate ^P^**c**^SS^, which is then related to the non-permuted vector **e** (Fig. 1b). The final max test then combines the row-based permutation test (using ^P^**e**) and the new column-based permutation test (using ^P^**c**^SS^) by only taking the largest P-value (least significant result), similarly as the previously described max test for the CWM approach without extension for ITV (Cormont et al. 2011, ter Braak et al. 2012).

**Fig. 1.**
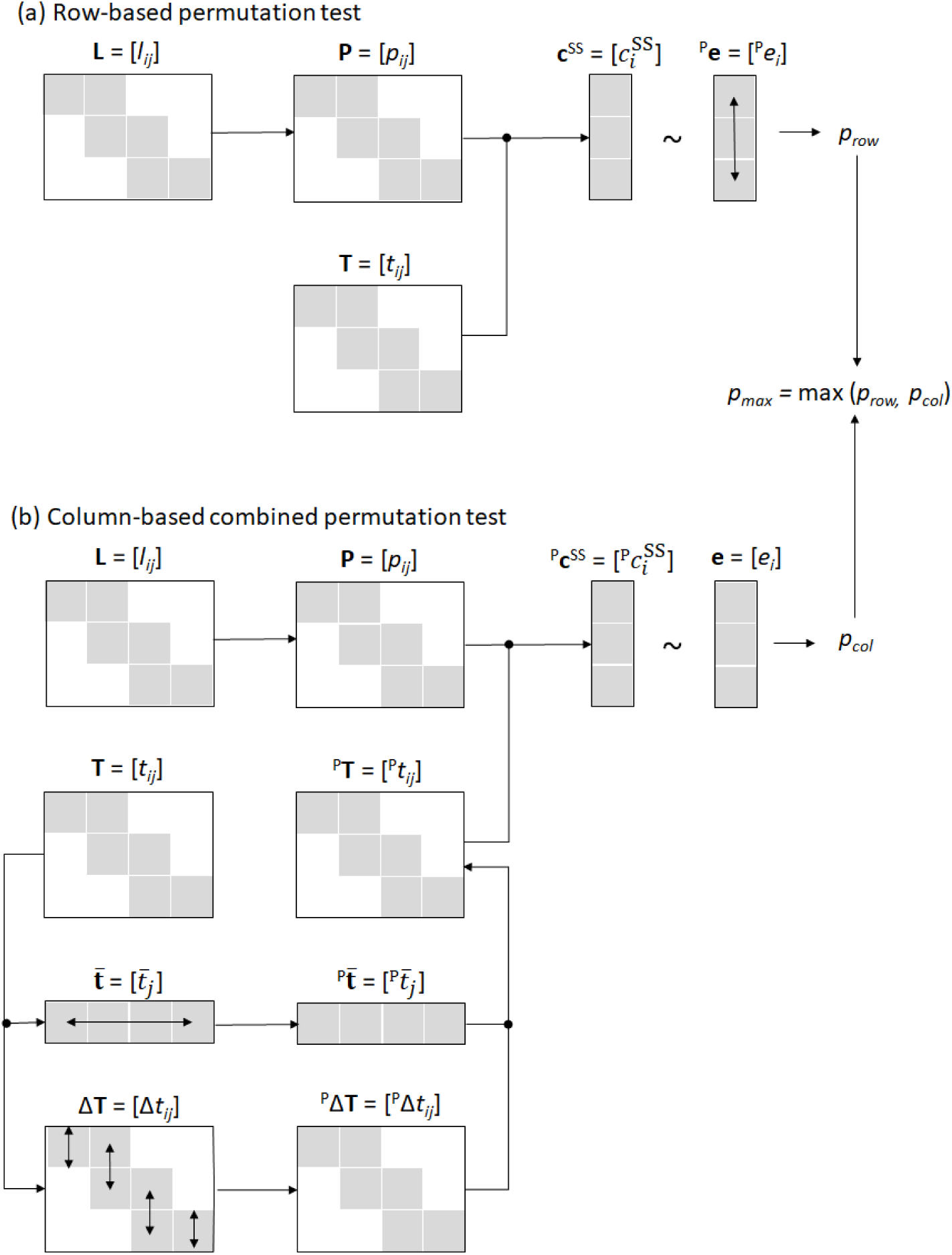
The schema of ITV-extended max test. (a) The row-based permutation test for site-specific trait values. (b) Combined column-based permutation test, with separate permutation of fixed trait values and intraspecific trait values. Max test combines *P-*values from both tests by selecting the higher one. Grey cells represent values of non-zero species abundances (in species composition matrix **L** and **P**) or values of traits and environmental variables which are not missing

We explored whether this newly proposed max test for the ITV-extended CWM approach can correctly control for the Type I error rate inflation for our 50 simulations, across different magnitudes of ITV. For this, we used the same 50 simulated datasets we used to quantify the inflation of the Type I error rate. For fixed CWM (*m* = 0), we replaced this newly modified column-based test with a test permuting only the mean trait values in (Fig.Fig. S1b). We repeated 1000 times each test for each combination of the dataset and ITV magnitude and plotted the average and standard deviation of the inflation index across the 50 simulations for each test against the ratio of intra- and interspecific trait variation, as described earlier.

We compared the test powers between the newly introduced ITV-extended max test for site-specific CWM-environment relationship with the standard parametric version of it and also with the standard parametric *F*-test for intraspecific CWM-environment relationship. We generated 1000 simulated community datasets for each combination of different sample sizes (10, 25, 50 and 100), different species numbers (10, 25, 50 and 100), and different magnitudes of ITV *m* (-1.0, -0.2, 0, 0.2, 1). For the community dataset of each combination of these parameters, we assigned trait values for each species in each site so that they match the site’s environmental conditions (while introducing a small level of stochastic noise). While interspecific trait values were always positively related to environmental variable, intraspecific variation was related either negatively (m < 0), not at all (m = 0) or positively (m > 0). We then calculated site-specific CWM and tested its relationship to the environment using the max and standard parametric tests. We also calculated intraspecific CWM and tested its relationship to the environment using the standard parametric *F*-test. For each combination of sample size, species number and ITV magnitudes, we reported the proportion of significant tests by calculating how many of the 1000 tests have been significant at P < 0.05 (the higher the proportion of significant tests, the higher the power of the test).

### Real-world dataset: leaf traits of woody species in the cloud forest of Taiwan

To illustrate the effect of ITV on community-level trait-environment relationships in a real-world dataset, we used data from the one-hectare vegetation plot in the cloud zone of northern Taiwan, hereafter termed the Lalashan Forest Dynamics Plot (see ESM Methods S1 for more details about locality and trait and environmental variables). We surveyed woody species in 25 systematically distributed 10 m × 10 m subplots. For broad-leaved species within each subplot, we measured leaf area (LA, mm^2^), specific leaf area (SLA, mm^2^/mg), leaf dry matter content (LDMC, mg/g) and leaf thickness (Lth, mm), following the protocols of Pérez-Harguindeguy et al. (2013). We measured leaf traits for 665 individuals of all 48 broad-leaf species. For each subplot, we also calculated a set of topographical parameters, including mean elevation (m), convexity (m) and windwardness. Additionally, we also calculated a hypothetical ’environmental’ factor that was directly calculated from the **L** matrix. This variable, the subplot scores on the first ordination axis of a correspondence analysis calculated on the **L** matrix (hereafter named CA1), presents the strongest possible predictor of subplot-level species composition since it is directly derived from it.

We used data from this case study for two subsequent analyses. In the first one, we evaluated the relationship between the Type I error inflation and the magnitude of ITV, for the four measured leaf traits and the two environmental factors, windwardness and CA1. We calculated Type I error rates for the relationships (linear regression, *F*-test) between the four site-specific and intraspecific CWMs, on the one hand, and windwardness and CA1, on the other hand (we did not consider the fixed CWMs in this analysis). Species for which no trait measurements were performed were removed from the **L** matrix. The two CWM vectors, **c**^SS^ and **c**^ITV^, were calculated as defined earlier. To break the relationship between traits and environment in this data, we permuted site-specific and intraspecific trait data and performed 10,000 independent permutations to quantify the inflation index, as described earlier. For each trait, we also calculated the relative ITV index and plotted the inflation index for each tested community-level trait–environment relationship against the relative ITV index. For site-specific CWMs, we additionally assessed if applying our newly introduced ITV-extended max test (with 999 permutations) could remove the Type I error inflation.

In the second analysis, we used the variation partitioning method introduced by Lepš et al. (2011) and modified by Fajardo & Siefert (2018) for linear regression to explore the specific community-level trait-environment relationships in our dataset and to quantify the effects of ITV, species turnover and their interaction on these relationships. We used all four measured leaf traits and the three measured topographical variables (but not CA1). For each trait and topographical variable combination, we calculated three linear regression models and tested them using the appropriate method: (i) **c**^F^ ∼ **e**, tested by the "original" max test, (ii) **c**^SS^ ∼ **e**, tested by the newly introduced ITV-extended max test, and (iii) **c**^ITV^ ∼ **e**, tested by the standard parametric *F*-test. For (i) and (ii), we also included standard parametric *F-*tests to allow comparison of the results with the correct test method (max test). We decided not to correct for multiple testing issue in this analysis, as we mainly compare differences between *P*-values calculated by standard parametric test and max permutation tests for individual trait-environment combinations. For each regression, we then calculated the sum of squares, where SS_fixed_ represents the effect of species turnover, SS_specific_ the total trait variation, and SS_ITV_ the effect of ITV, respectively. We subsequently partitioned those variations by the formula SS_specific_ = SS_fixed_ + SS_ITV_ + covariation. All sum of square values were then rescaled to a percentage scale, where SS_specific_ was set to 100%.

All calculations were performed in R 4.3.3. (R Core Team 2024). The R code is provided as Supplementary File 2, and the real-world data set is at https://doi.org/10.5281/zenodo.10936367. The simulated datasets were generated using the *simcom* package (Zelený, version 0.1.0, https://github.com/zdealveindy/simcom), max permutation tests for the fixed CWM-environment relationships were performed with the *weimea* package (Zelený, version 0.1.18, https://github.com/zdealveindy/weimea), correspondence analysis with the *vegan* package (Oksanen et al., version 2.5-7) and the partitioning of among-plot trait variation with functions modified from the *cati* package (Taudiere & Violle, 2015, version 0.99.3).

## Results

### Simulated community data

The relationships between site-specific CWMs and the environment, tested using standard parametric tests, have an inflated Type I error rate, where the inflation is negatively related to the magnitude of ITV (Fig. 2a). The Type I error inflation is highest at the smallest relative ITV index, approaching the inflation for fixed CWM. The inflation seems almost absent if the relative ITV index is higher than 3, and there is no apparent inflation when ITV is unconstrained (*m* = ∞). The relationship between intraspecific CWMs and environment, tested using standard parametric tests, does not have an inflated Type I error rate (Fig. 2b), and consequently shows no relationship with the relative ITV index. This test, however, seems rather conservative, with all inflation index values below one.

**Fig. 2.**
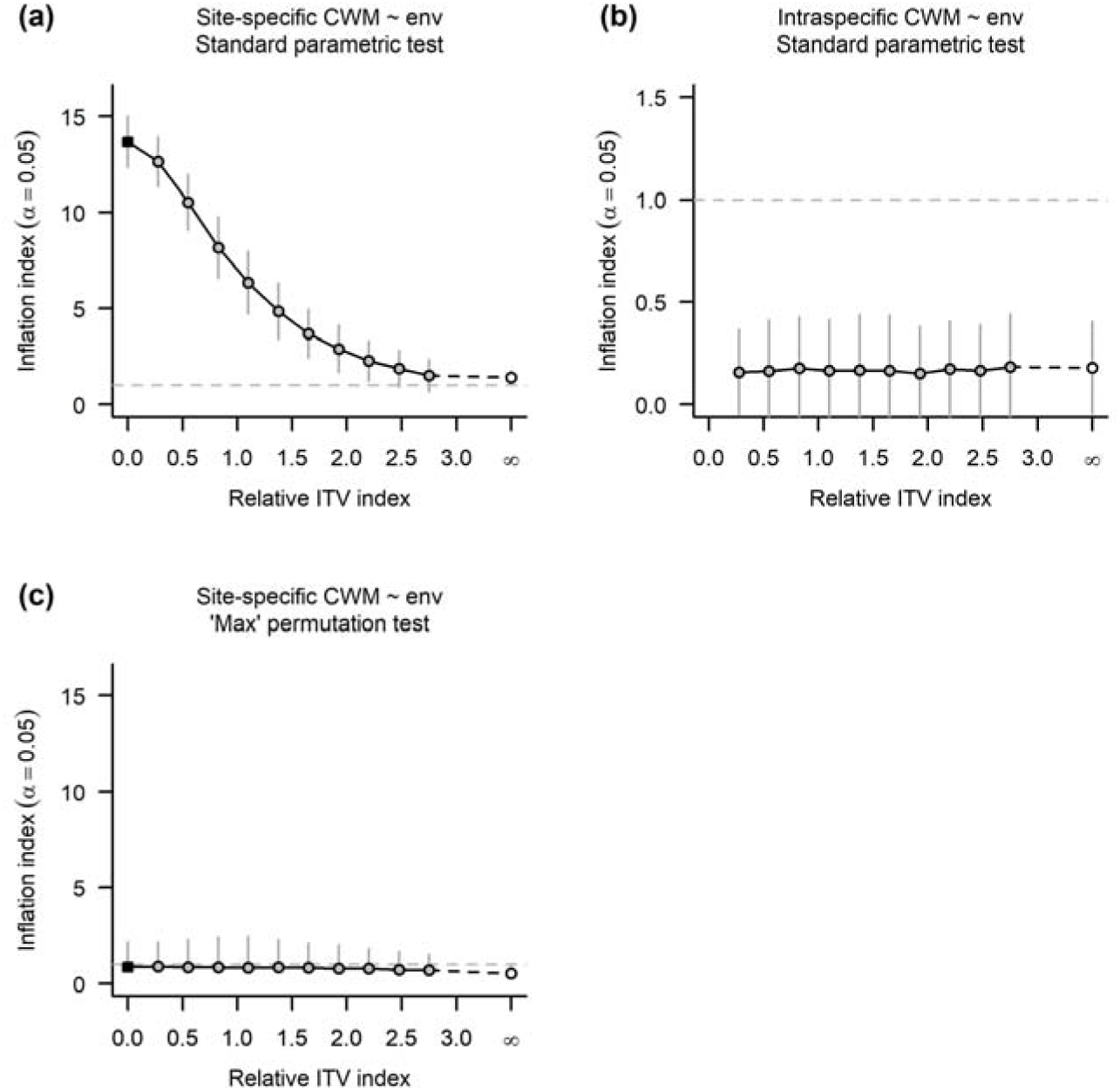
The effect of the magnitude of intraspecific trait variation (relative ITV index) on inflation index (mean + standard deviation) of the linear regression between (a, c) site-specific CWM or (b) intraspecific CWM and the ’environmental variable’ of the simulated data, tested by (a, b) standard parametric test and (c) "max" permutation test. Black square = fixed CWM (cf. site-specific CWM with no ITV). Relative ITV index equal to infinity (∞) represents situation when site-specific trait matrix with trait values randomly sampled from normal distribution was used. No inflation occurs if the inflation index is equal or lower to 1 (indicated by dashed horizontal line)

Our newly introduced ITV-extended max test successfully controls for the Type I error rate inflation of site-specific CWMs for all magnitudes of ITV (Fig. 2c). For fixed CWMs (ITV = 0), this test reverts to the "original" max test, which also controls for the Type I error rate inflation (Fig. 2c). The relationship of intraspecific CWMs and environment was not inflated when tested by standard parametric *F*-tests, so no permutation-based correction was necessary.

When comparing the power of individual tests, the newly introduced ITV-extended max test of site-specific CWM-environment relationship shows consistently lower power than the parametric version, especially for datasets with a lower number of sites and species (ESM Fig. S3). The power of the max test also sharply decreases for datasets with negative values of ITV magnitude, where intraspecific trait variation is negatively related to the environment, while interspecific trait variation is positively related to the environment (power of the parametric test remains relatively high). Standard parametric F-test for intraspecific CWM-environment relationship has consistently lower power in smaller datasets and lower absolute values of ITV magnitude.

### Leaf traits of woody species in the cloud forest

Regression of site-specific CWMs against the two environmental variables (the measured windwardness and generated CA1) showed an inflated Type I error rate for all four measured traits (Fig. 3a). The inflation index values were, overall, higher for regressions against CA1 compared to windwardness. While the inflation index showed a somewhat decreasing trend with increasing relative ITV for CA1, no relationship was observed for windwardness. All regressions of intraspecific CWMs against CA1 and windwardness had an inflation index close to 1, with no apparent trend along the increasing relative ITV index (Fig. 3b). The newly introduced ITV-extended max test also successfully addressed the Type I error inflation in this dataset (inflation index close to or lower than 1).

**Fig. 3.**
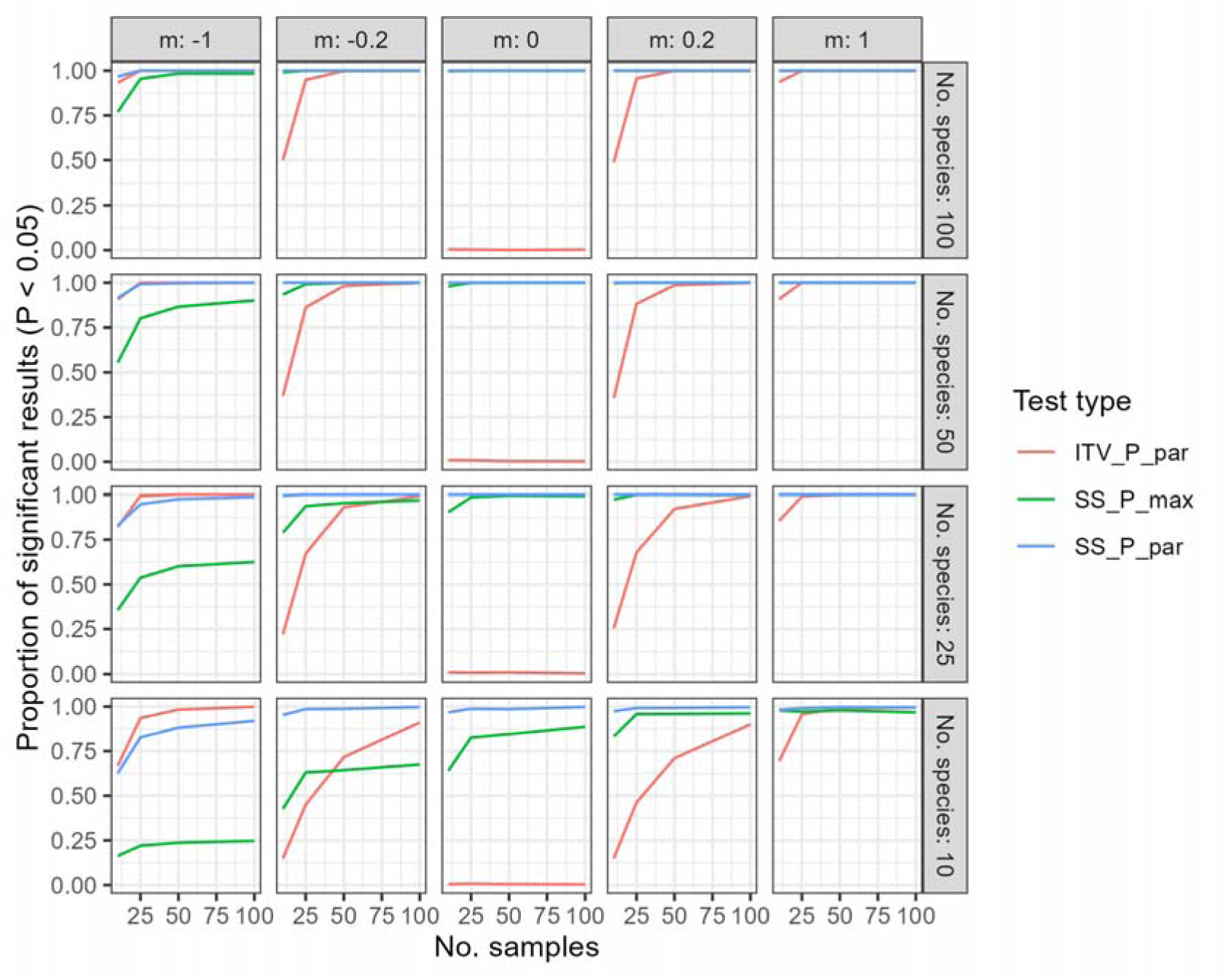
Power analysis comparing ITV-extended max test of site-specific CWM-environment relationship (SS_P_max) with its standard parametric version (SS_P_par), and also with standard parametric F-test for intraspecific CWM-environment relationship (ITV_P_par). For each combination of number of samples (No. samples), number of species (No. species) and magnitude of ITV (m), 1000 compositional datasets were calculated, and the proportion of tests significant at P < 0.05 was counted. Interspecific trait variation is always positively related to environmental variable, while intraspecific is negatively related for negative *m* and positively for positive *m*

**Fig. 4.**
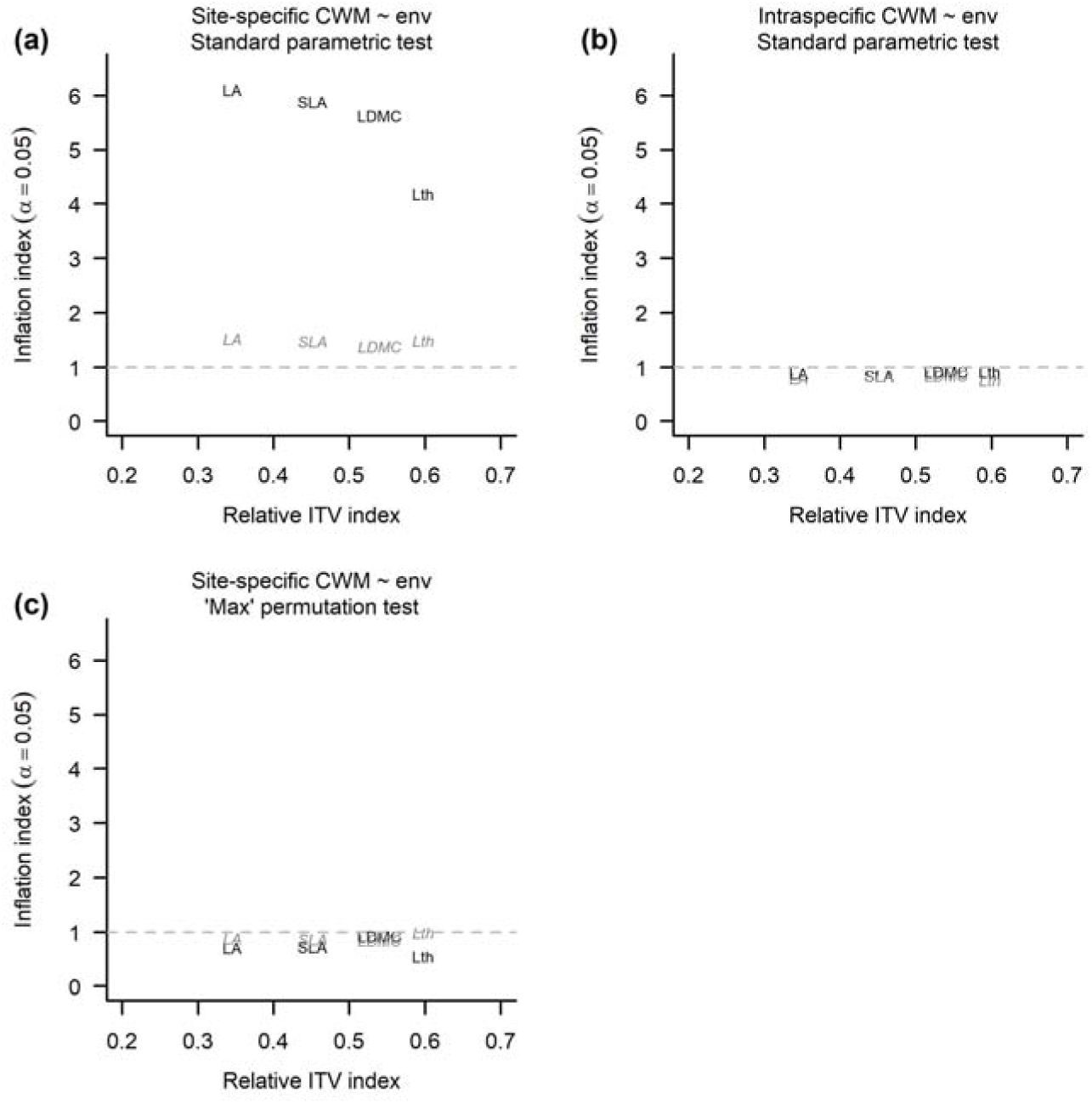
The effect of the magnitude of intraspecific trait variation (relative ITV index) on inflation index (mean + standard deviation) of the linear regression between (a, c) site-specific CWM or (b) intraspecific CWM and windwardness (gray italics) and CA1 (black) of the cloud forest data, tested by (a,b) standard parametric test and (c) "max" permutation test. No inflation occurs if the inflation index is equal or lower to 1 (indicated by dashed horizontal line). LA = leaf area, SLA = specific leaf area, LDMC = leaf dry matter content, Lth = leaf thickness

From the four site-specific and fixed CWM-environment relationships in our dataset, which were significant (P < 0.05) when tested by F-test, two became insignificant following max test correction (ESM Tab. S1). For site-specific CWMs, Lth was positively and SLA negatively related to windwardness (ESM Fig. S4b and c, respectively) based on both the parametric and max tests (ESM Tab. S1). For fixed CWMs, LA was positively related to elevation and SLA negatively related to convexity when tested with parametric tests, but based on the max test, both relationships are only marginally significant (P < 0.1) (ESM Tab. S1). Finally, for the intraspecific CWMs (tested only by parametric test), we found a negative relationship with windwardness for both LA and SLA (ESM Fig. S4a&c) and a positive relationship between Lth and windwardness (ESM Fig. S4b). None of the three CWMs for LDMC were significantly related to any measured environmental variable. Variance partitioning of the trait–environment relationship into the effect of species turnover, ITV and their covariation showed a considerable positive covariation fraction for the SLA and Lth relationships with windwardness (ESM Fig. S5).

## Discussion

We illustrate with both simulated and real community data that testing community-level trait-environment relationships suffer from inflated Type I error rate when CWMs include (among-site) ITV in a similar way as when CWMs are calculated from fixed species-level trait values. We also showed that the extent of this inflation decreases with increasing amounts of ITV; for very low ITV magnitudes, it approaches the inflation of trait– environment relationships using fixed CWMs, while for high ITV magnitudes, it is almost non-existent (Fig. 2a). At the range of ITV magnitude observed in our real-world dataset (0.35-0.60), inflation remains strong. The newly introduced max test extended for ITV proved to control Type I error rate for the full range of simulated ITV magnitude, and we suggest to use it whenever exploring site-specific CWM -environment relationships. Indeed, our simulation dataset suggests that levels of ITV need to be more than three times the amount of interspecific trait variation before inflation becomes neglectable, so that our new max test is likely to be needed in most real data studies.

Our real-world dataset is rather small concerning both the number of species (48) and sites (25), and is also quite homogeneous in terms of environmental conditions (due to the small spatial extent). Consequently, we would expect the magnitude of ITV in our study (0.35-0.60, or 35-60%, respectively) to be less extensive than for more species-rich communities across strong environmental gradients at large spatial scales. Surprisingly, other studies have nonetheless found lower ITV magnitudes, both at local (15-33%, Jung et al. 2010) and global scales for plant communities (32%, Siefert et al. 2015). It seems therefore unlikely that the extent of ITV in real datasets would be sufficiently high (>300%) to overcome the Type I error inflation for site-specific CWM-environment relationships without using a max test correction, at least for plant communities. A detailed review of community-level studies, including ITV, might be helpful to quantify the range of ITV magnitudes for different taxa, environmental strengths and spatial scales. Also note that the comparison of ITV magnitude with other studies is slightly hampered by the use of several alternative measures for ITV magnitude (cf. Lepš et al. 2006, Albert et al. 2010a, Albert et al. 2010b, de Bello et al. 2011, Siefert et al. 2015).

The use of trait values measured at the level of individual sites ("site-specific ITV"), as in this study, is just one example of how ITV can be incorporated into CWM-based trait– environment relationships. ITV covers any type of intraspecific trait variation, from variation among leaves of a single tree to variation among individuals of a species occurring on different continents. Specifically for CWM-based trait–environment relationships, the amount of included ITV can gradually range from the inclusion of only ’habitat-specific’ or ’region-specific’ ITV, where sites of one habitat or region are characterised by fixed species-level trait values (e.g. Lepš et al. 2011, Helsen et al. 2018), to the inclusion of fine-scale intra-site ITV (trait variation among individuals in a site) (e.g. Carlucci et al. 2015). The severity of the Type I error inflation is expected to decrease along this gradient since the more detailed ITV information is included, the more likely ITV magnitude will be considerable. Our study nonetheless suggests that the Type I error corrections will remain necessary for any study where the magnitude of ITV is lower than 3.

The actual level of Type I error inflation in real-world datasets is also influenced by several other parameters next to the amount of ITV. As shown using a similar simulation model used in our study, inflation increases with decreasing beta diversity of the species compositional data, increasing the number of community samples used in the analysis and increasing strength of the link between the environmental variable and the species composition data (Zelený 2018). The strength of the **e**–**L** link, in particular, likely explains why the inflation is high for site-specific CWM related to CA1 (which is intrinsically strongly linked to the **L** matrix) and low when related to the real measured environmental variable, windwardness (which has a much weaker link to matrix **L**). The quantitative change in P-values after applying the newly introduced ITV-extended max test could be rather low (as in our real-world dataset, where two results became not significant at P < 0.05, but still staying marginally significant at P < 0.1). This may feel as if the max test is actually unnecessary; however, unless applied, one will not know the true extent of the Type I error inflation. We suggest that researchers apply the new test routinely to make sure that the results they obtain are fair and do not suffer from an inflated Type I error rate.

Surprisingly, the relationships between the intraspecific CWMs and the environment showed no Type I error inflation when tested with standard parametric tests. Even more, the inflation rates of this test are consistently lower than 1, in both simulated data and real-world data, and the power of this test is lower in datasets of smaller sample sizes and lower species numbers. We hypothesise that the lack of power is caused by the way intraspecific CWMs are calculated: the matrix of site-specific trait values is converted into the matrix of intraspecific trait values by centring the species’ traits, resulting in intraspecific trait values and CWM’s which tend to have values close to zero, and thus very low variance. In the case of the simulated dataset, which is less noisy, this behaviour is quite pronounced, resulting in low inflation index values (<0.4), while for the noisier real-world data, this behaviour is less pronounced, with inflation index values only slightly below 1. Detailed analyses should be performed in the future to uncover whether this is the reason for low power of this test.

Our study demonstrates the problem of Type I error inflation for site-specific CWMs – environment relationships assessed specifically using linear regression. However, we assume the same problem applies to other methods assessing trait-environment relationships with the CWM approach, including correlation (parametric or non-parametric), weighted regression (ter Braak et al. 2018a) or ANOVA. We assume that our new ITV-extended max test can solve this inflation in all of these methods since they belong (or are closely related) to the same statistical family of general linear models. As shown by ter Braak et al. (2018a) for fixed CWMs, the "original" max test also applies to this whole range of methods. It nonetheless remains useful to formally evaluate the sensitivity of these different methods to the Type I error rate inflation and their respective power.

In our analysis of the trait-environment relationship on real-world cloud forest data, we deliberately ignored the Type I error inflation problem associated with multiple testing, which arises when conclusions are based on the results of several (non-independent) tests on the same dataset. Note that this issue is independent of the Type I error rate inflation explored in this study. When the CWM approach is used to identify multiple trait-environment relationships, an additional correction of significance levels for this multiple testing issue is necessary to avoid inflated family-wise Type I error rates (see Wright 1992). We suggest basing this correction on the number of trait-environment pairs, not on the overall number of tests performed; i.e. no matter whether the study focuses on only a single CWM type (e.g. the site-specific one) or all three CWMs, each P-value should be adjusted by the value calculated as the number of traits × the number of environmental variables. The correction for multiple testing will also require a higher number of permutations for each individual test to allow the adjusted *P*-values to reach values lower than the selected significance threshold (e.g. α = 0.05). In case of a higher number of traits and environmental variables, the adjustment for multiple testing will result in low overall power of the test. A solution to this may be applying multivariate CWM-RDA analysis (Nygaard & Ejrnaes, 2004) on all traits and environmental variables at once, with a single global test evaluating an overall trait-environmental relationship (separately for individual CWM types).

### Practical considerations

For researchers using the CWM approach extended for ITV, we suggest the following workflow. First, make sure that the hypothesis you are testing is the one for which the CWM-environment relationship has inflated Type I error (consult alternative options described in Zelený, 2018, and Lepš & de Bello, 2023). If yes, then it is important to know that from the three CWMs calculated within the CWM approach extended for ITV, namely fixed, site-specific and intraspecific CWM, only the first two are prone to Type I error inflation. For fixed CWMs, we suggest using the original max (permutation) test, as introduced by Peres-Neto et al. (2017), which is currently available in the *weimea* R package (Zelený, unpublished; https://github.com/zdealveindy/weimea). For site-specific CWMs, the max test extended for ITV, as introduced in this study, can be used by applying the custom-made functions provided in the R code accompanying this manuscript (https://github.com/zdealveindy/ITV_CWM). For intraspecific CWMs, standard parametric tests do not suffer from Type I error rate inflation, and no correction is needed.

When using the results of previously published studies that applied the CWM approach extended for ITV without controlling for the Type I error inflation, be aware that some of them may be overly optimistic. As previously shown for the CWM approach without extension for ITV, the Type I error inflation in fixed CWM-environment relationships correlates positively with dataset size and strength of the link between species composition and environment, and negatively with the overall species beta diversity (Zelený 2018). We show that for assessing site-specific CWM-environment relationships using the CWM approach extended for ITV, the Type I error inflation is additionally negatively dependent on the magnitude of ITV.

## Acknowledgements

Our thanks go out to all members of the Vegetation Ecology Lab at the National Taiwan University and several volunteers who participated in the vegetation data collection campaign of the Lalashan Forest Dynamics Plot, especially Ting Chen and Kun-Sung Wu.

## Declarations

### Ethics approval

Not applicable.

### Consent to participate

Not applicable.

### Consent for publication

Not applicable.

### Availability of data and material

The real-world dataset is on GitHub (https://doi.org/10.5281/zenodo.10936367).

### Code availability

The R code used for analysis is included in Supplementary File 2.

### Authors’ contributions

DZ conceived the idea, analysed data, and led manuscript writing. KH contributed to discussions and improved the manuscript. YNL collected and analysed real-world datasets and commented on the manuscript. The authors declare no conflicts of interest.

