## Supplement for "Extending the CWM approach to intraspecific trait variation: how to deal with overly optimistic standard tests?"

Electronic Supplementary Material (ESM)

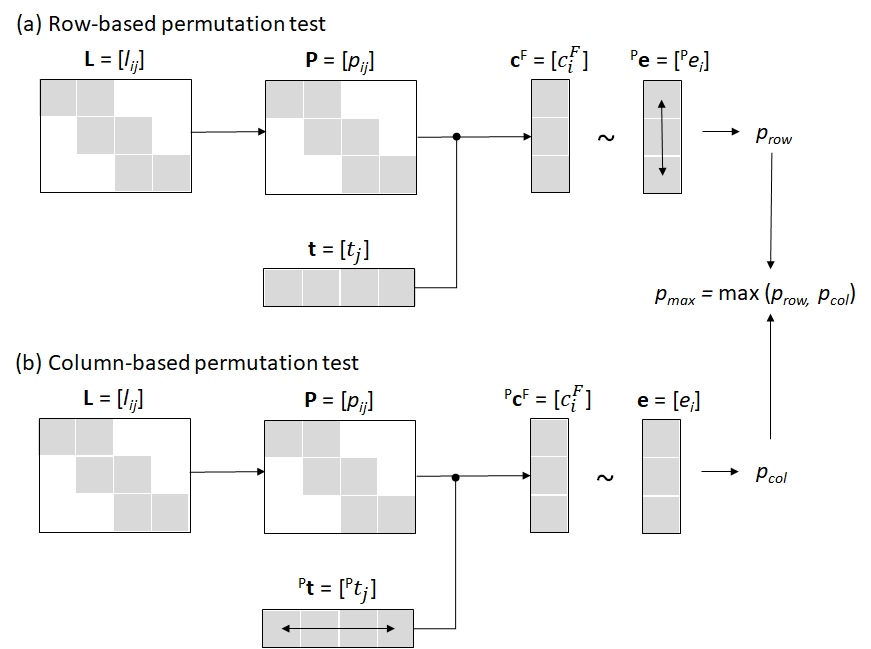

**Figure S1.** The schema of (a) the row-based and (b) the column-based permutation test for fixed trait values CWM. The resulting *P-*values from both tests are combined into the max test. Grey cells represent values of non-zero species abundances (in species composition matrix **L** and **P**) or values of traits and environmental variables that are not missing.

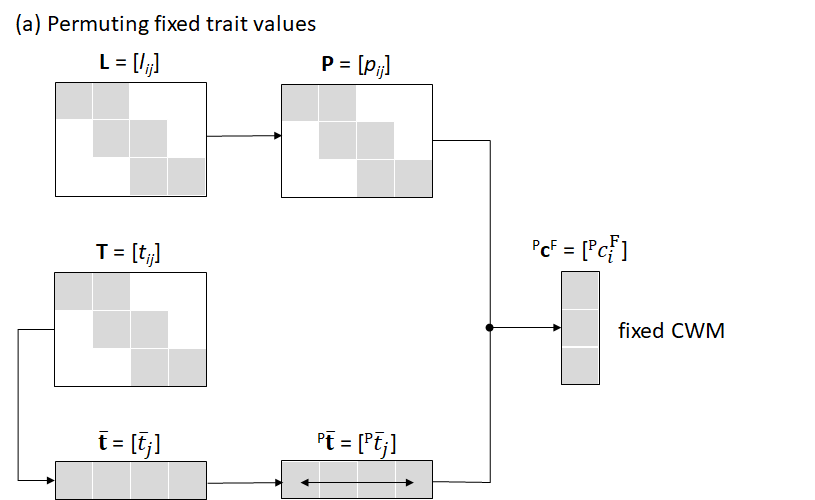

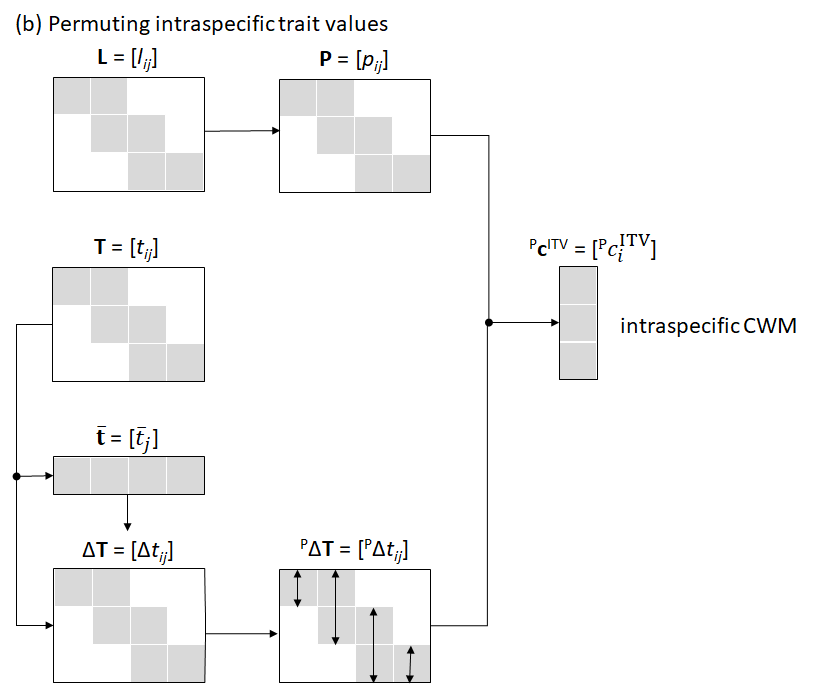

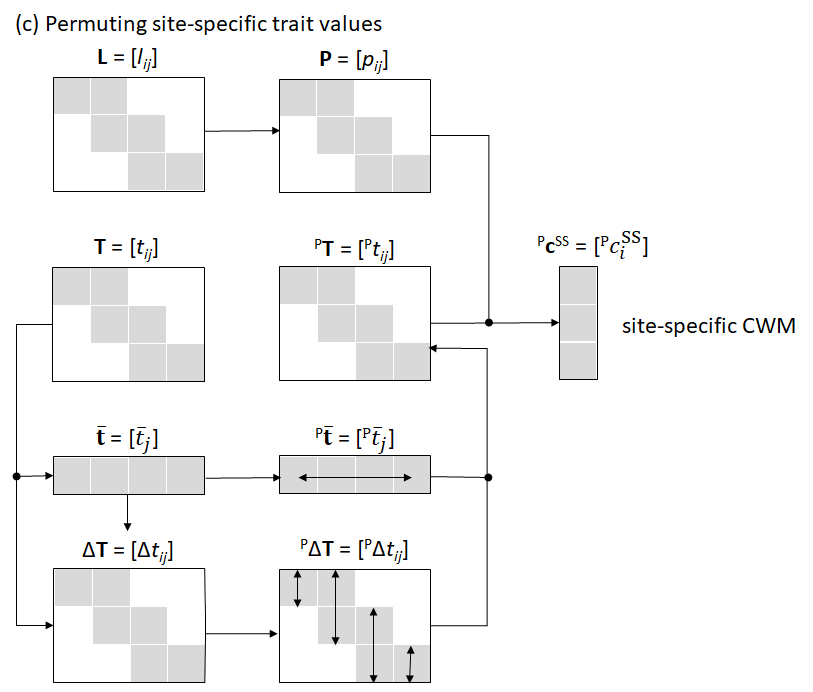

**Figure S2.** The schema of calculating permuted (a) fixed CWM, (b) intraspecific CWM, and (c) site-specific CWM. Horizontal or vertical arrows within the vectors and matrices indicate the direction of the permutation. Grey cells represent values of non-zero species abundances (in species composition matrix **L** and **P**) or values of traits that are not missing (in trait vectors or matrices).

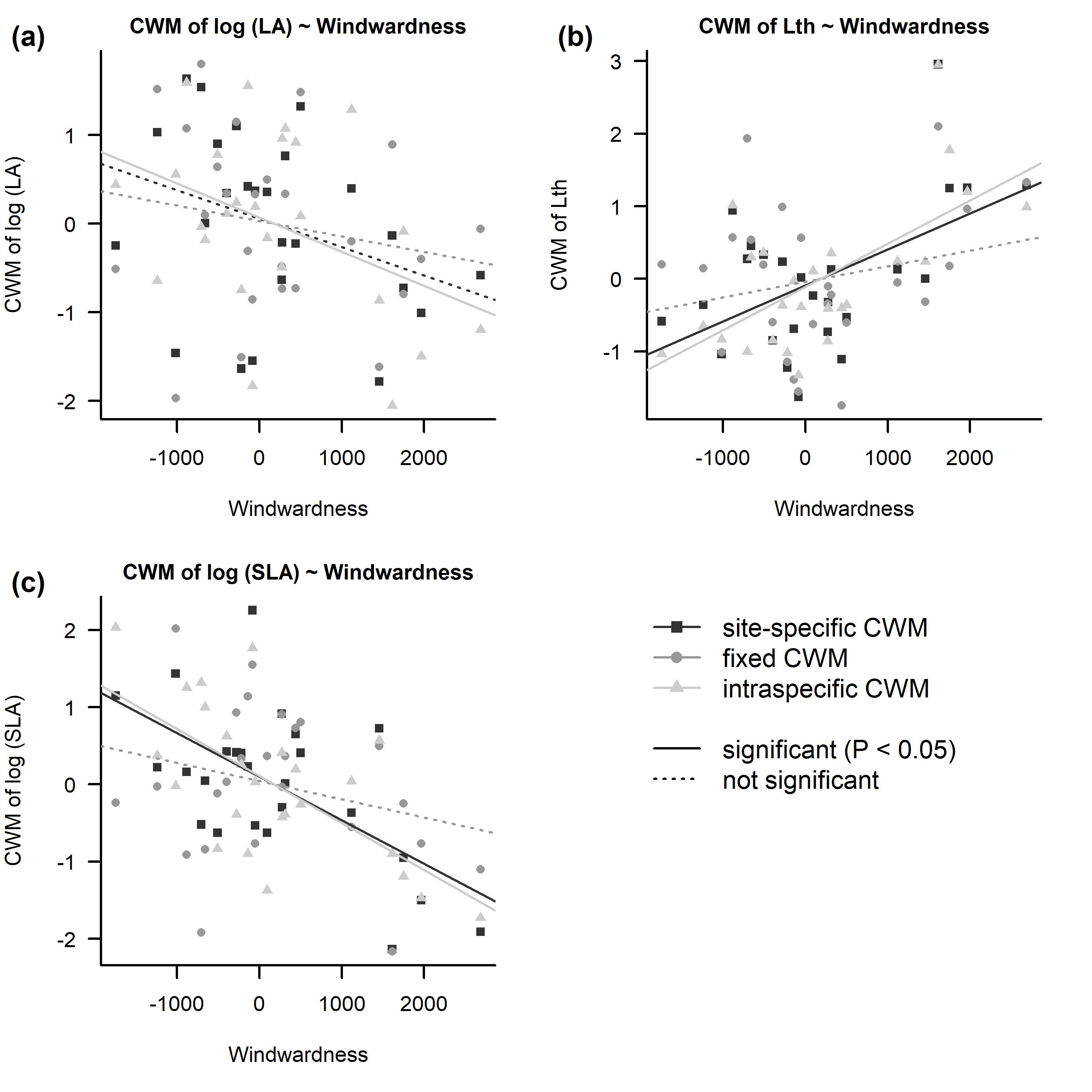

**Figure S3.** Regression between windwardness and the site-specific, fixed, and intraspecific CWM of (a) leaf area, (b) leaf thickness, and (c) specific leaf area. All CWMs were z-transformed. Regressions significant at *P* < 0.05 (max test in the case of site-specific and fixed CWM, parametric test in the case of intraspecific CWM) were visualized by a solid regression line, non-significant regressions by a dashed line.

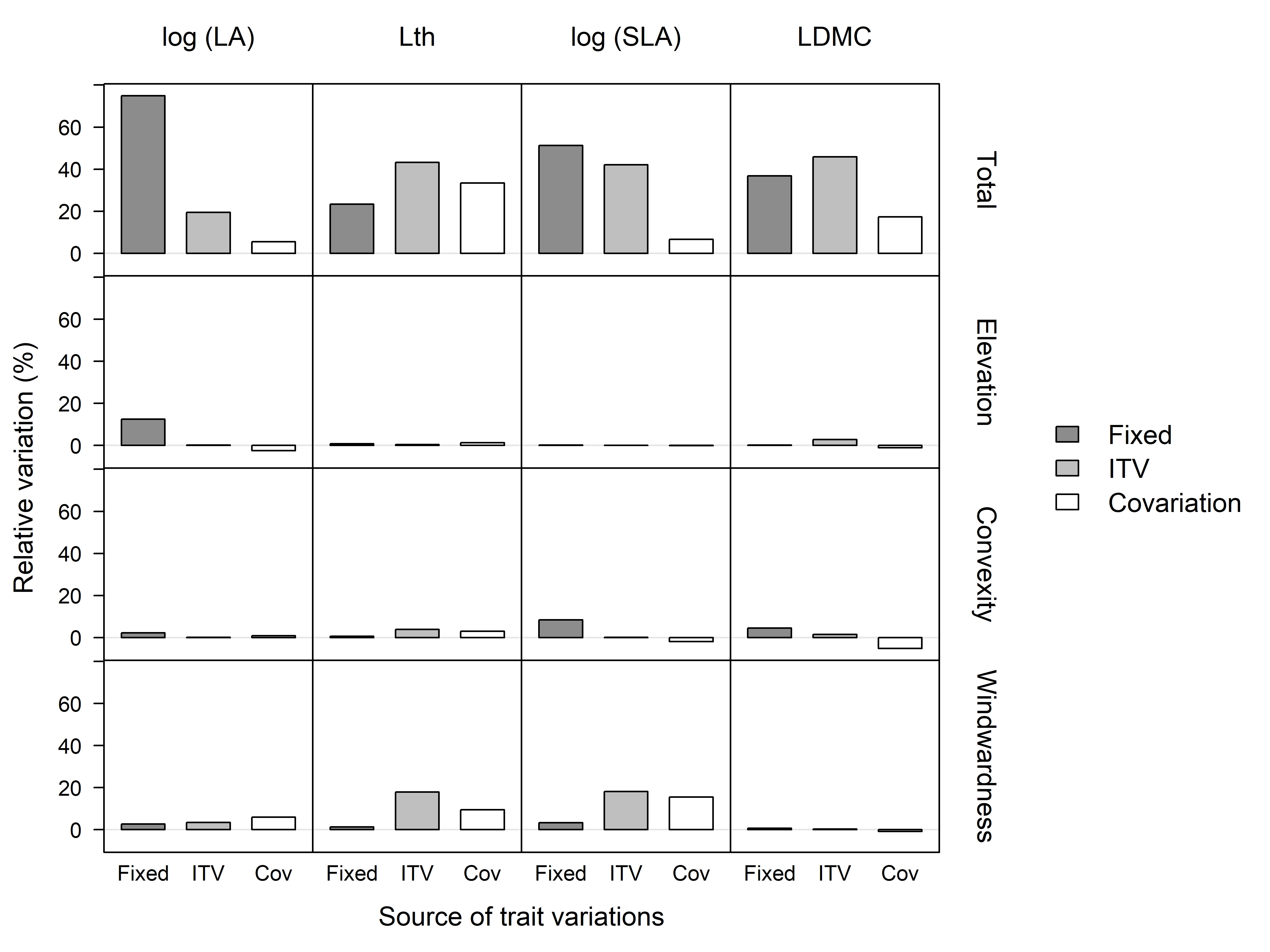

**Figure S4.** Partitioning of variation in community-level trait responses to environmental variables in the cloud forest of northern Taiwan. The total among-subplot variance in leaf traits explained by environment was partitioned into the effect of species turnover (fixed CWM), ITV (intraspecific CWM) and covariation. Partitioning was done for four traits: leaf area (LA), leaf thickness (Lth), specific leaf area (SLA), and leaf dry matter content (LDMC), and three topographical variables: elevation, convexity and windwardness.

**Table S1.** Leaf traits of woody species in the cloud forest: regression of site-specific (SS), fixed (F) and intraspecific (ITV) CWM on environmental variables, tested by parametric *F*-test (*P*_par_) and max permutation test (*P*_max_; ITV-extended max test was used for site-specific CWM, and “original” max test for fixed CWM). Significant results (*P* < 0.05) are in bold.

| Environmental variable | CWM type | LA |  |  |  |  | Lth |  |  |  |  | SLA |  |  |  |  | LDMC |  |  |  |
| --- | --- | --- | --- | --- | --- | --- | --- | --- | --- | --- | --- | --- | --- | --- | --- | --- | --- | --- | --- | --- |
|  |  | *r*^2^ | *F* | *P*_par_ | *P*_max_ |  | *r*^2^ | *F* | *P*_par_ | *P*_max_ |  | *r*^2^ | *F* | *P*_par_ | *P*_max_ |  | *r*^2^ | *F* | *P*_par_ | *P*_max_ |
| elevation | SS | 0.101 | 2.58 | 0.122 | 0.179 |  | 0.025 | 0.60 | 0.448 | 0.518 |  | 0.001 | 0.01 | 0.911 | 0.923 |  | 0.018 | 0.42 | 0.523 | 0.628 |
|  | F | 0.166 | 4.58 | **0.043** | 0.095 |  | 0.035 | 0.85 | 0.367 | 0.497 |  | 0.002 | 0.05 | 0.822 | 0.881 |  | 0.003 | 0.07 | 0.799 | 0.860 |
|  | ITV | 0.006 | 0.15 | 0.704 |  |  | 0.011 | 0.25 | 0.624 |  |  | 0.000 | 0.01 | 0.940 |  |  | 0.061 | 1.48 | 0.236 |  |
| convexity | SS | 0.031 | 0.73 | 0.403 | 0.442 |  | 0.073 | 1.82 | 0.190 | 0.218 |  | 0.066 | 1.63 | 0.215 | 0.252 |  | 0.008 | 0.18 | 0.678 | 0.691 |
|  | F | 0.030 | 0.70 | 0.411 | 0.486 |  | 0.024 | 0.56 | 0.461 | 0.549 |  | 0.164 | 4.50 | **0.045** | 0.091 |  | 0.120 | 3.14 | 0.089 | 0.154 |
|  | ITV | 0.004 | 0.08 | 0.778 |  |  | 0.089 | 2.25 | 0.147 |  |  | 0.003 | 0.06 | 0.810 |  |  | 0.033 | 0.79 | 0.384 |  |
| windwardness | SS | 0.120 | 3.13 | 0.090 | 0.104 |  | 0.285 | 9.18 | **0.006** | **0.007** |  | 0.370 | 13.50 | **0.001** | **0.003** |  | 0.001 | 0.02 | 0.890 | 0.909 |
|  | F | 0.035 | 0.84 | 0.369 | 0.488 |  | 0.054 | 1.31 | 0.265 | 0.364 |  | 0.065 | 1.60 | 0.219 | 0.311 |  | 0.018 | 0.43 | 0.520 | 0.609 |
|  | ITV | 0.172 | 4.79 | **0.039** |  |  | 0.412 | 16.13 | **0.001** |  |  | 0.431 | 17.41 | **0.000** |  |  | 0.006 | 0.14 | 0.710 |  |

### **Methods S1**

#### Simulated community data with ITV (species composition and environment)

To generate simulated compositional data structured by an environmental variable (vector **e**) we used the COMPAS model proposed by Minchin (1987) and extended by Fridley et al. (2007). The extended model allows the creation of a simulated community by generating *S* unimodal species response curves along a vector **e** of fixed length, where each species response curve (represented by a Beta function, Minchin 1987) quantifies the probability with which a random individual found at a given gradient location is assigned to that given species. The species composition of individual sites is then generated by randomly selecting *n* locations along the environmental gradient, and assigning a predefined number of individuals to different species at each site, according to the species (response-curve-defined) occurrence probability at that site. In our simulation, we set the number of species *S* = 50, the number of sites *n* = 25, and 100 individuals were sampled in each site. The width of each species response curve (one of the parameters of the Beta function) is generated as a random number from a uniform distribution between 2500 and the total length of the environmental gradient, 5000 units. Sites along the gradient were sampled between 500 and 4500 units to avoid gradient edges with a lower density of species response curves. The simulation model returns two objects: a vector **e** (positions of sites along the gradient), and a *n*-by-*S* composition matrix **L** (with numbers in cells representing the counts of individuals of a given species in a given site). This simulation was performed 50 times with the same number of species and sites, resulting in 50 independent sets of **e** and **L**. See the Materials & Methods section of the main manuscript for a description of how a matrix of site-specific trait values (**T**) was generated using this data.

#### Real-world dataset: leaf traits of woody species in the cloud forest of Taiwan

To illustrate the effect of ITV on community-level trait-environment relationships in a real-world dataset, we used data from the one-hectare vegetation plot in the cloud zone of northern Taiwan (24°42′25″ N, 121° 26′29″ E, 1758-1782 m a.s.l.), hereafter termed the Lalashan Forest Dynamics Plot. The plot is located on a wide mountain ridge, with several dry gullies and a windward slope in the eastern part of the plot. The vegetation is defined as *Chamaecyparis* montane mixed cloud forest (Li et al. 2013), with coniferous cypress *Chamacyparis obtusa* var. *formosana* dominating the canopy, and several evergreen broad-leaf tree species dominating the subcanopy (e.g., *Neolitsea accuminatissima, Quercus sessilifolia*, *Rhododendron formosanum, Trochodendron aralioides*). Within the 100 m × 100 m Lalashan Forest Dynamics Plot, established following ForestGEO protocol (Condit 1988), we surveyed woody species in 25 systematically distributed 10 m × 10 m subplots. We recorded the diameter at breast height (DBH) and species identity of all woody individuals with a DBH ≥ 1 cm, and used the relative number of individuals and relative basal area (derived from DBH) to calculate the importance value index (IVI) per subplot for each species (Curtis 1959). IVI values were then organized in the subplot × species **L** matrix. In total, we surveyed 1110 individuals of 49 species (including 48 broadleaved and one coniferous species).

For 1–3 individuals of each broad-leaved species randomly chosen within each subplot, we collected three mature leaves for trait measurements. For each leaf, we measured leaf area (LA, mm^2^), specific leaf area (SLA, mm^2^/mg), leaf dry matter content (LDMC, mg/g), and leaf thickness (Lth, mm), following the protocols of Pérez-Harguindeguy et al. (2013). Leaf-level trait values were first averaged per individual and subsequently per species in each subplot, to obtain a species × site-specific trait values **T** matrix. Since the distribution of site-specific trait values of LA and SLA was strongly right-skewed, we log_10_-transformed them before further analysis. We measured leaf traits for 665 individuals of all 48 broadleaved species.

For each subplot, we also calculated a set of topographical parameters, including mean elevation (m), convexity (m), and windwardness. The mean elevation of the subplot and convexity were calculated from the elevation of corner piles, following Valencia et al. (2004). Windwardness is a combination of aspect and slope, expressed as ‘easterness’ × slope, where ‘easterness’ is the aspect folded along the east-west axis, rescaled into +90° for the E and -90° for the W direction. Windwardness is expected to be related to the effect of the chronic north-eastern (winter) monsoon winds. We additionally calculated a hypothetical ‘environmental’ factor, directly calculated from the **L** matrix and presenting the strongest possible predictor of subplot-level species composition. This variable was calculated as the subplot scores on the first ordination axis of a correspondence analysis calculated on the **L** matrix (hereafter named CA1).
